# Generation of liver organoids from human induced pluripotent stem cells as liver fibrosis and steatosis models

**DOI:** 10.1101/2021.06.29.450347

**Authors:** Hoi Ying Tsang, Paulisally Hau Yi Lo, Kenneth Ka Ho Lee

**Affiliations:** MOE Key Laboratory for Regenerative Medicine, School of Biomedical Sciences, Chinese University of Hong Kong, Shatin, Hong Kong; (H.Y.T.); (P.H.Y.L.)

## Abstract

**Background & Aims:** Liver cirrhosis is a major cause of death worldwide, and its prevalence is growing rapidly due to the growth of obesity and diabetes population with non-alcoholic fatty liver disease (NAFLD). Yet, no effective therapeutics have been developed to treat NAFLD or its more advanced stage, non-alcoholic steatohepatitis (NASH). This has raised great concern for a representative liver model to be developed so that novel drugs could be screened, identified and developed. Presently, we aim to develop a liver organoid entirely from human induced pluripotent stem cells (hiPSC) to model liver fibrogenesis and NAFLD.

**Methods:** Hepatoblasts (HBs), mesenchymal stem cells (MSCs), hepatic stellate cell (HSCs) and endothelial cells (ECs) were derived from hiPSCs, allowed to self-organized and differentiated into liver organoids. Liver functions, transcriptomic and protein expression of liver organoids were characterized and validated. Liver organoids were exposed to thioacetamide (TAA) and free fatty acids (FFA) to be induced into liver disease model.

**Results:** The liver organoids we fabricated were highly vascularized, exhibited liver-specific functions and hepatic cellular spatial organization. The presence of liver specific ECs, macrophages and cholangiocytes were found within our organoids. TAA induced fibrosis in our liver organoids that exhibited diminished liver functions, elevated pro-inflammatory cytokines and fibrosis-related gene expression, as well as extensive collagen deposit. Organoids treated with FFA developed steatosis, inflammation and fibrosis.

**Conclusions:** We generated a novel method, that is Matrigel-independent and size-controllable, for making human liver organoids. These organoids can potentially be utilized as tissue-mimetic *in vitro* model for high throughput screening to identify drugs that can be used to treat liver fibrosis and NAFLD.

## INTRODUCTION

Liver fibrosis is a wound healing response, when extracellular matrix (ECM) is over-secreted upon liver injuries or chronic insults, such as virus, drugs, alcohol and excess lipids. Persistence liver injury leads to chronic inflammation, ECM accumulation, extensive scarring and ultimately cirrhosis, which is a major cause of death worldwide (1). Non-alcoholic fatty liver disease (NAFLD) affects about 20% of the world population and shows rapidly increasing prevalence due to the growing obesity and diabetes population (2). Nevertheless, no drug has yet been approved as a treatment to NAFLD. The failure in drug discovery is attributed to the high failure rate in clinical trials, caused by the low concordance between *in vitro* and *in vivo* data, as well as between animals and human studies (3, 4). A more representative *in vitro* model that includes functional human liver cells is therefore essential to bridge the gap between preclinical and clinical studies.

Development of liver fibrosis and non-alcoholic steatohepatitis (NASH), an advanced type of NAFLD, involves the interplays between liver parenchymal cells (PCs), i.e. hepatocytes, and non-parenchymal cells (NPCs) including Kupffer cells (residing liver macrophages), liver sinusoidal endothelial cells (LSECs) and importantly hepatic stellate cells (HSCs) (5). HSCs can be activated from quiescent vitamin storing cells into ECM secreting cells by danger associated molecular patterns (DAMPs) released from damaged hepatocytes, and are considered a key player in fibrogenesis (6).

Human induced pluripotent stem cells (hiPSCs) can be induced to differentiate into various liver associated cell types, to provide a promising and reliable alternative to primary hepatocytes for drug screening. Protocols have now been developed to differentiate hiPSCs into liver PCs (7–9) and NPCs (10–14), enabling the development of personalized medicine. To improve the maturation of iPSC-derived hepatocyte-like cells (HLC) and achieve a more tissue-mimetic *in vitro* model, different approaches have been developed. For example, HLCs have been cultured in 3D manner and co-cultured with NPCs and were found to restore the cell-cell and cell-ECM interactions as seen in native liver (15).

Liver organoids derived from iPSCs is another approach for procuring an organotypic model. Current protocols mainly involves self-organization and hepatic differentiation of iPSC or using iPSC-derived endoderm cells in Matrigel (16–20). These organoids display many liver functions, but most of these organoids contained only epithelial cell type, hepatocytes and cholangiocytes, but not other cell types normally found in the liver. The presence of NPC-like cells have been reported in some organoid models, but formed as a result of uncontrolled spontaneous differentiation that is highly varied between organoids. The use of Matrigel also diminishes the control of organoid size which creates great inter-batch variations which compromise the potential use of organoids in high throughput drug screening. Takebe’s group demonstrated the formation of liver bud by mixing iPSC-derived hepatic endoderm cells, mesenchymal stem cells and human umbilical vein endothelial cells (HUVECs) on a soft matrix that mimics liver organogenesis (21). The liver bud was vascularized and differentiated into mature liver tissues after *in vivo* transplantation, but its potential use as an *in vitro* model for liver fibrosis and NAFLD has not been evaluated.

In this study, we have generated a novel method of making vascularized liver organoids, using iPSC-derived hepatoblasts (HBs), mesenchymal stem cells (MSCs), hepatic stellate cells (HSCs) and endothelial cells (ECs). We were able to precisely control the size of the organoids and the spatial distribution of the various cell type within the organoids. Moreover, our liver organoid model exhibited liver-specific functions and architecture and has the ability to recapitulate complex events associated with liver fibrosis and steatosis.

## MATERIALS AND METHODS

### iPSCs culture and differentiation

hiPSC cell line (iBU-2, generated from reprogrammed dermal fibroblasts by retrovirus carrying the Yamanaka gene factors) was maintain on Matrigel-coated culture dishes in Essential 8 Flex Medium (ThermoFisher Scientific). The protocols for culturing cell lines and differentiating the hiPSCs into HBs, MSCs, HSCs and ECs are described in the supplementary materials.

### Differentiation of liver organoids

iPSC-derived HBs (7 days post-differentiation), iPSC-derived MSCs (cell passage 6 - 10), iPSC-derived HSCs (cell passage 2 - 4) and iPSC-derived ECs (cell passage 2 - 4) were trypsinized and seeded into ultra-low attachment 96-well plate (BRANDplates® inertGrade) at 5000 cells in 200 µl media. The culture medium was composed of 1:1 mixture of EGM-2 and HCM without EGF supplemented with 2µM A83-01, 1 mM dexamethasone and 100 nM Dihexa. The medium was used at a ratio of 10:7.2:2:0.8 for HBs, EC, MSC and HSC. Supplements 10µM Y-27632 and 10ng/ml FGF2 were added to the differentiation medium on day 1-2 and day 1-4, respectively. 75% of medium was replaced with fresh medium 3 times a week for 14 days.

### Liver function tests

ELISA for human albumin: Culture media were collected from organoid cultures and stored at -80°C until use. Quantification of secreted albumin was performed using a Human Albumin Quantification Kit (Bethyl Laboratories). Procedure according to manufacturer’s instructions and absorbance was measured using a microplate spectrophotometer at 450nm. Indocyanine green (ICG) uptake and excretion: Organoids were incubated in 1 mg/ml ICG for 1 hour, washed with PBS three times and then cultured in fresh media for 4 hours to allow excretion of ICG.

### Real-time quantitative PCR (qPCR)

Total RNA was extracted using RNAiso Plus reagent (TaKaRa) with or with a Direct-zol RNA microprep kit (Zymo Research) according to the manufacturer’s instructions and stored at -80°C. Concentration and purity of RNA was determined with Nanodrop 2000 Spectrophotometer (ThermoFisher) and reverse transcription of 2 μg of RNA was performed using PrimeScript RT Master Mix (TaKaRa). RT-qPCR was performed with SYBR Premix Ex Taq ((Tli RNaseH Plus), TaKaRa) on QuantStudio 7 Flex Real-Time PCR System (Applied Biosystems). qPCR primers used were listed in supplementary methods.

### Whole-mount immunofluorescent staining

The organoids were washed with PBS twice, fixed in 4% Paraformaldehyde (PFA) for 60 mins with mild shaking at 4°C, and then temporarily stored in PBS at 4°C. Before staining, the organoids were permeabilized in 0.2% Tween-20 and 0.5% Triton-X100 prepared in PBS for 15 minutes twice and blocked with 5% BSA in 0.2% Triton-X100-PBS for 3 hours at room temperature. The organoids were then incubated in primary antibodies diluted with blocking solution overnight at 4°C with mild shaking. Organoids were washed with 1% BSA in 0.1% Triton-X100-PBS for 2 hours at 4°C thrice, before incubation in secondary antibodies diluted in blocking solution and 2 μg/ml DAPI overnight at 4°C. Organoids were washed for 2 hours thrice, and cleared overnight at 4°C in 2.5M, 60% glycerol solution before imaging with confocal microscope (Leica SP8). Antibodies and the dilutions used were listed in supplementary methods.

### Histological study

The organoids, after 60 mins fixation in 4% PFA, were washed with PBS. They were then embedded in 2% agarose/2.5% gelatin gel and fixed in 10% neutral buffered formalin overnight at 4°C. The organoids in the gel were dehydrated, embedded in paraffin blocks and sectioned at 4 μm using a microtome (Leica). The sections were used for hematoxylin-eosin (H&E), periodic acid-Schiff (PAS) and pico-sirius red (PS Red) staining.

### Thioacetamide (TAA) and free fatty acid (FFA) treatment

TAA was dissolved in culture media, oleic acid (Sigma) and palmitic acid (Sigma) were conjugated with fatty acid-free BSA at molar ratio of 6:1. Organoids were exposed to TAA (10/25 mM) or FFA-BSA conjugate (oleic acid and palmitic acid mixed at 1:1 molar ratio) (200/400/800 μM) for 7 days, before the collection of media and organoids for analyses. Cell viability of organoids was measured with CellTiter-Glo 3D Cell Viability Assay (Promega) according to manufacturer’s instructions.

### Graphs and statistical analysis

Graphs were constructed using a GraphPad Prism 9 software and data presented as average value from three or more independent experiments with standard deviation as error bars. The data were subjected 2-tailed Student *t* test between two groups. P-values were shown in graphs as asterisks, with * representing p <0.05, ** representing p< 0.01 and “*** representing p<0.001. P-value<0.05 was regarded as statistically significant.

## RESULTS

### Differentiation of iPSCs into hepatic parenchymal and non-parenchymal cell types

We first differentiated hiPSCs into liver PCs and NPCs. To validate the hepatic differentiation potentials of our hiPSC line, the cells were differentiated using a three-step protocol (9) first to definitive endoderms (DE) on day 2, then hepatoblasts (HB) on day 7 and finally into hepatocyte-like cells (HLC) on Day 21 (Fig. S1). Differentiation efficiency was determined by flow cytometric analysis of hepatic markers, with 62.89 ± 5.03% iPSC-derived HBs expressing AFP and 58.90 ± 8.04% expressing HNF4A (Fig. S1C). The iPSC-derived HLCs that we created were then profiled and showed that they expressed hepatic markers AFP, ALB and HNF4A (Fig. 1A and 1B). The HLCs could also secrete albumin and urea, and were capable of storing of glycogen and lipid droplets (Fig. 1C and 1D). We also generated iPSC-derived MSCs that expressed MSC markers CD29, CD44, CD75, CD90 and CD105 with the absence of haematopoietic marker CD34 (Fig. 1E and 1F). These MSCs possessed tri-lineage differentiation potential to become osteoblasts, adipocyte and chondrocytes (Fig. 1G). We used FACS to purify CD31^+^ cells from iPSC-derived ECs. These cells exhibited a “cobblestone” morphology that is typical for endothelial cells (Fig. S3C) and maintained CD31 and VECAD expressions for more than 4 passages with a small portion of CD34^+^ cells (Fig. 1H and 1I). The ECs were able to form extensive tubular networks when plated onto Matrigel (Fig. 1J). The iPSC-derived HSCs showed a star-like morphology (Fig. S3F) and expressed HSC markers ALCAM, HGF, PDGFRB, ICAM1, COL1 and aSMA (Fig. 1K and 1L). These HSCs were able to store retinoid droplets that gave them an autofluorescence appearance under UV (Fig. 1M). Moreover, the HSCs could be activated by TGFB1 to induce an increase in cell proliferation, expression of stress fibres and activated HSC markers (Fig. 1N), as well as loss of retinoid droplets (Fig. S3J).

**Fig. 1.**
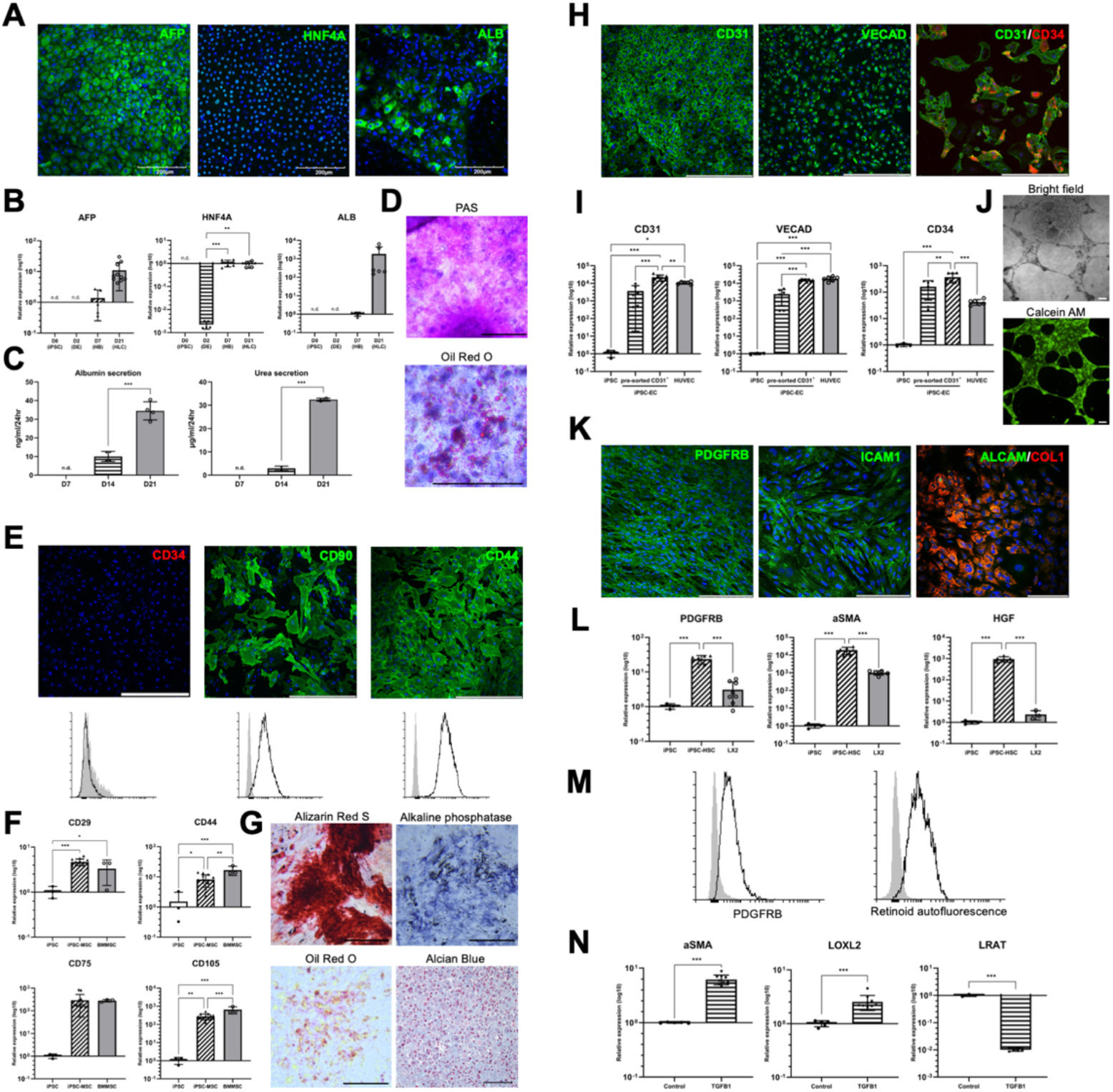
Differentiation of hiPSCs. (A) Immunofluorescent staining of hepatocyte markers (AFP, ALB and HNF4A) in D21 hepatocyte-like cells (HLCs). (B) Gene expression at Day (D)2, D7 and D21 after differentiation as determined by RT-qPCR, with GAPDH as internal control. (C) Albumin and urea secretion of HLCs at D14 and D21 HLCs. (D) PAS and Oil Red O staining in HLCs. (E) Representative immunofluorescent staining image and flow cytometric analysis of haematopoeitic marker (CD34) and MSC markers (CD44, CD90) in iPSC-derived MSCs. IgG controls displayed in grey. (F) Gene expression of MSC markers (CD29, CD44, CD75 and CD105) in iPSC-derived MSCs assessed by RT-qPCR. (G) Alkaline phosphatase and Alizarin Red S staining for osteoblasts, Oil Red O staining for adipocytes and Alcian Blue staining for chondrocytes that were differentiated from iPSC-MSCs. (H) Immunofluorescent staining for EC markers (CD31, VECAD and CD34) in CD31^+^ iPSC-derived ECs. (I) Gene expression iPSC-derived ECs before and after FACS assessed by RT-qPCR. (J) Tube formation of iPSC-derived ECs on matrigel. (K) Immunofluorescent of HSC markers (PDGFRB, ICAM1 and ALCAM/COL1) in iPSC-derived HSCs. (L) Representative flow cytometric analysis of iPSC-derived HSCs positive for HSC marker PDGFRB and auto-fluorescent retinoid droplets. Negative controls (PDGFRB: IgG; Retinoid: culture media without retinol) displayed in grey. (M) Gene expression of HSC markers (HGF, PDGFRB and aSMA) in iPSC-derived HSCs assessed by RT-qPCR. (N) expression of activation markers (aSMA, LOXL2) and quiescent marker (LRAT) genes in iPSC-HSCs after 72-hour 10ng/ml TGFB1 exposure by RT-qPCR. Results are shown as mean ± SEM (n≥3). *p <0.05; **p <0.01; ***p <0.001 using 2-tailed student’s *t* test. Scale bar = 200 μm.

### Optimization of initial cell number for organoid differentiation

We co-cultured day 7 iPSC-derived HBs with iPSC-derived NPCs (composing of MSCs, ECs and HSCs) for 14 days with starting seed number of 3000, 4000, 5000, 6000, 7000 and 8000 cells to form iPSC-derived HB/NPC organoids. The size of the HB/NPC organoids, cell viability (quantified by ATP content), and ability to secrete albumin were increased according to the initial seeding cell number (Fig. 2A and 2B). For seeding cell number over 5000, the organoids showed stagnated level of albumin secretion with large variation between samples. Therefore, we determined that 5000 cells was the optimal seeding number for constructing our HB/NPC organoids which allowed production of organoids with the highest consistency and high level of albumin secretion.

**Fig. 2.**
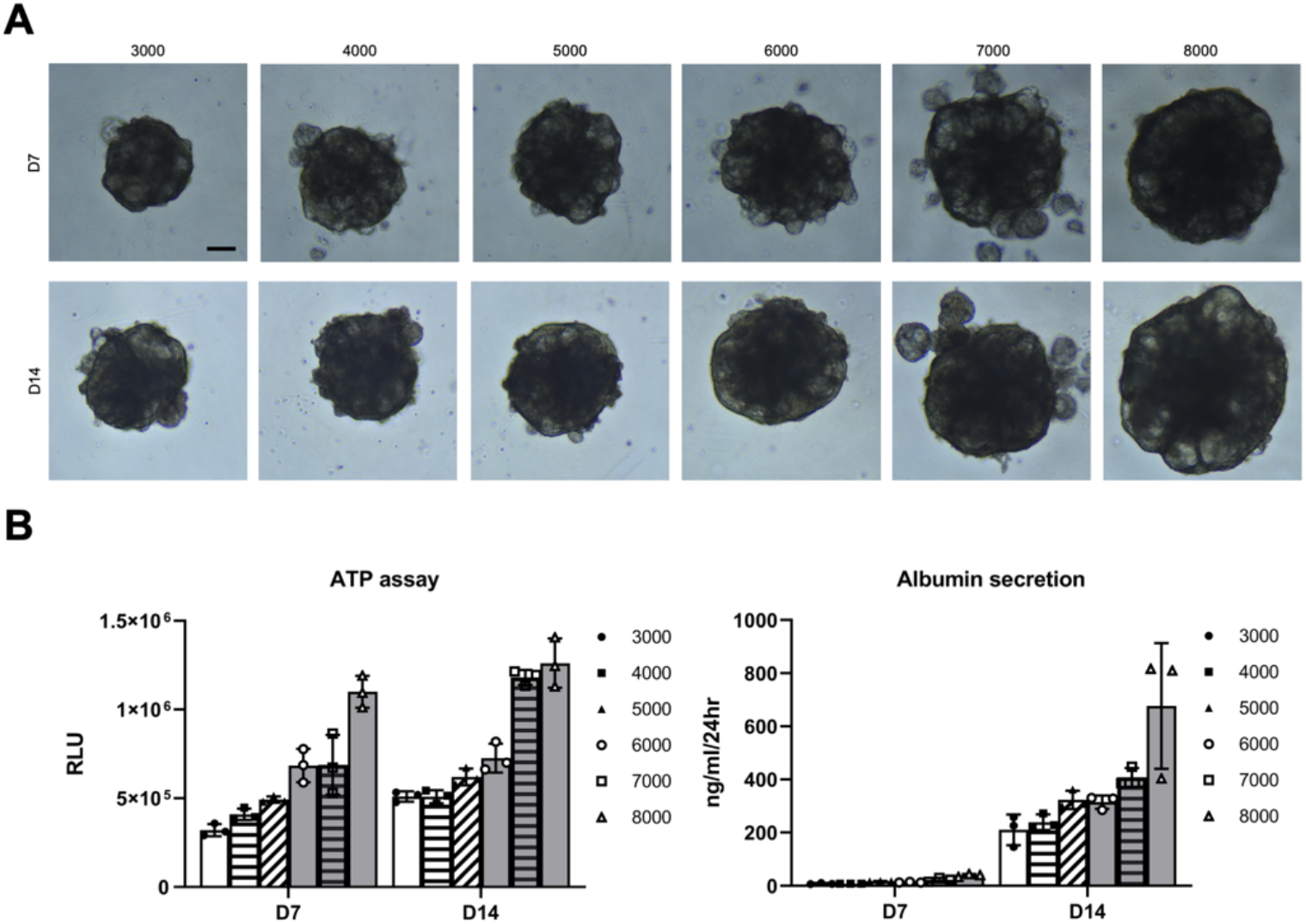
Effects of initial cell seeding number for organoid formation. (A) Morphology of liver organoids at different seeding number on day 7 and 14. Scale bar = 200 μm. (B) Cell viability measured using ATP bioluminescence assay and albumin secretion measured by ELISA of organoids on day 7 and 14 at different seeding number (n=3).

### Liver organoid formation improves differentiation of functional hepatocytes

We investigated whether the formation of liver organoids enhanced the differentiation and maturation of HBs into functional hepatocytes. As controls, we co-cultured iPSC-derived HBs and NPCs in 2D culture, and also created organoids composed of only iPSC-derived HBs. These organoids formed with iPSC-derived HBs were characterized (Fig. S5), then compared with the iPSC-derived HB/NPC organoids. iPSC-derived HB/NPC organoids showed significantly higher expression level of hepatocyte markers ALB, AAT, TAT, bile canaliculi marker MRP2, glucose metabolizing enzymes G6PC, and cytochrome P450 (CYP) enzymes 2B6, 2C9 and 2C19 than iPSC-derived HB organoids and 2D HB/NPC co-culture (Fig. 3A). The iPSC-derived HB organoids secreted albumin at 9.95 ± 1.09 ng/ml/24hr and 62.68 ± 11.02 ng/ml/24hr on day 7 and day 14 respectively, while iPSC-derived HB/NPC organoids secreted 13.10 ± 4.53 ng/ml/24hr and 366.73 ± 111.57 ng/ml/24hr, suggesting a significantly improved liver function if the organoids incorporated NPCs (Fig. 3B).

**Fig. 3.**
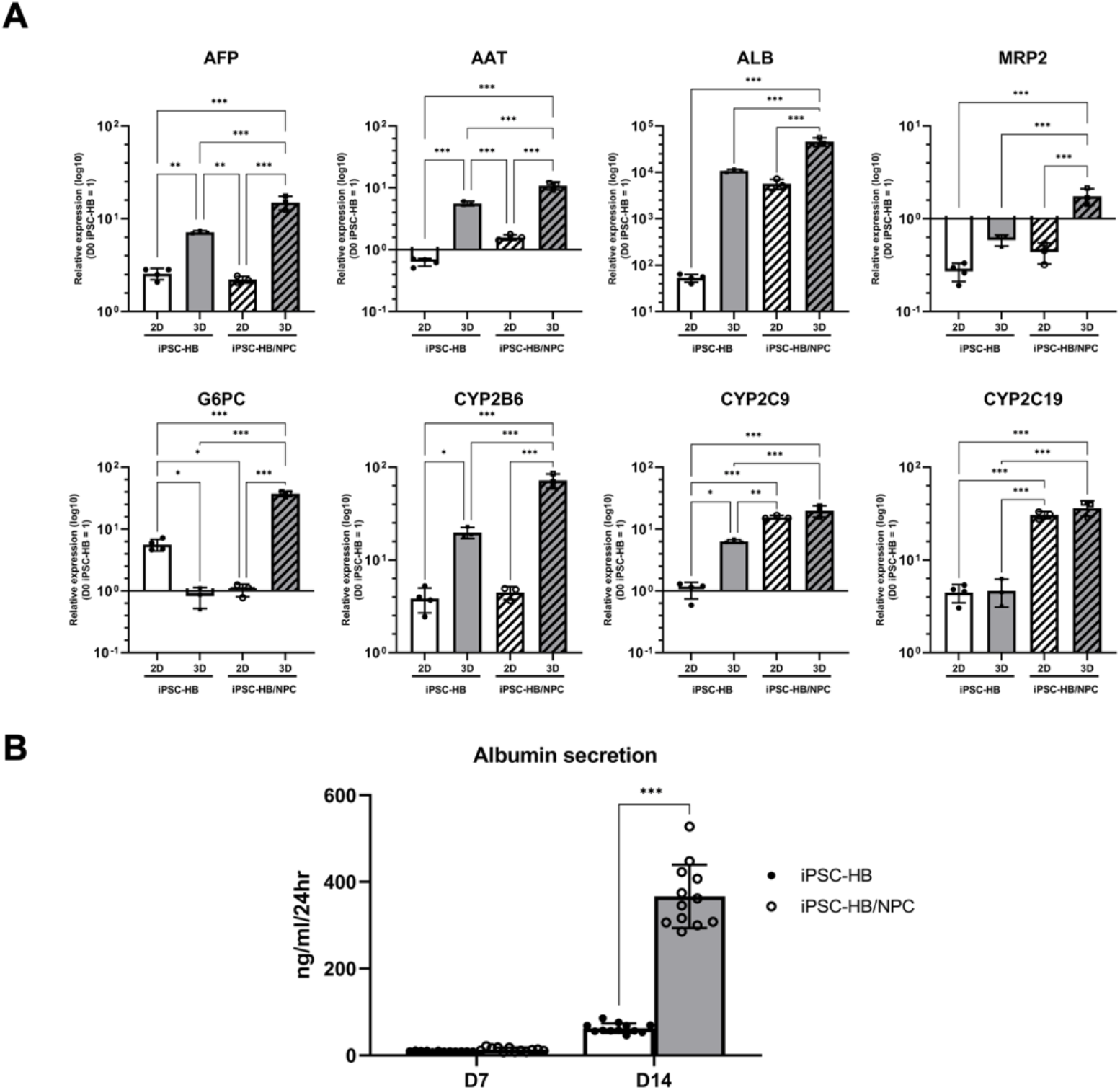
3D co-culture of iPSC-derived HBs and NPCs enhanced hepatic differentiation. RT-qPCR showing ALB, AAT, TAT, MRP2, G6PC, CYP2B6, 2C9 and 2C19 expressions at day 14, in 2D and 3D iPSC-derived HB mono-culture, 2D and 3D iPSC-derived HB and NPC co-culture. (B) Albumin secretion by iPSC-derived HB only organoids and iPSC-derived HB/NPC organoids on day 14. Results are shown as mean ± SEM (n≥3). *p <0.05; **p <0.01; ***p <0.001 using 2-tailed student’s *t* test.

### Liver organoids exhibit liver-specific functions

We investigated CYP enzyme expression in liver organoids following treatment with CYP enzymes inducer Rifampicin. RT-qPCR analysis revealed that organoids treated 25 μM Rifampicin for 72 hours were induced to express CYP2B6, 2C19 and 3A4 at 1.50 ± 0.20, 2.75 ± 0.41, and 3.14 ± 0.53 folds respectively, suggesting an inducible drug metabolism in our liver organoids (Fig. 4A). Immunofluorescent staining confirmed the presence of cells that expressed ALB and AAT, mainly locating at the outer surface of organoids (Fig. 4B). This outer layer of cells were positive for epithelial marker ECAD and expressed liver zonation markers CPS1 and GS, indicating the identity of these cells as mature hepatocytes (22). Polarization of hepatocytes was validated by the expression of tight junction marker ZO1 and MRP2. Bile canaliculi network was visualized by CDCFDA efflux assay (Fig. 4C). We determined that Indocyanine green dye was rapidly absorbed within 1 hour and excreted out within 4 hours, after removal of the dye from the liver organoids (Fig. 4C). (23). Histological sections of the liver organoid were positively stained for PAS, further confirming the hepatic functions of our organoids (Fig. 4C).

**Fig. 4.**
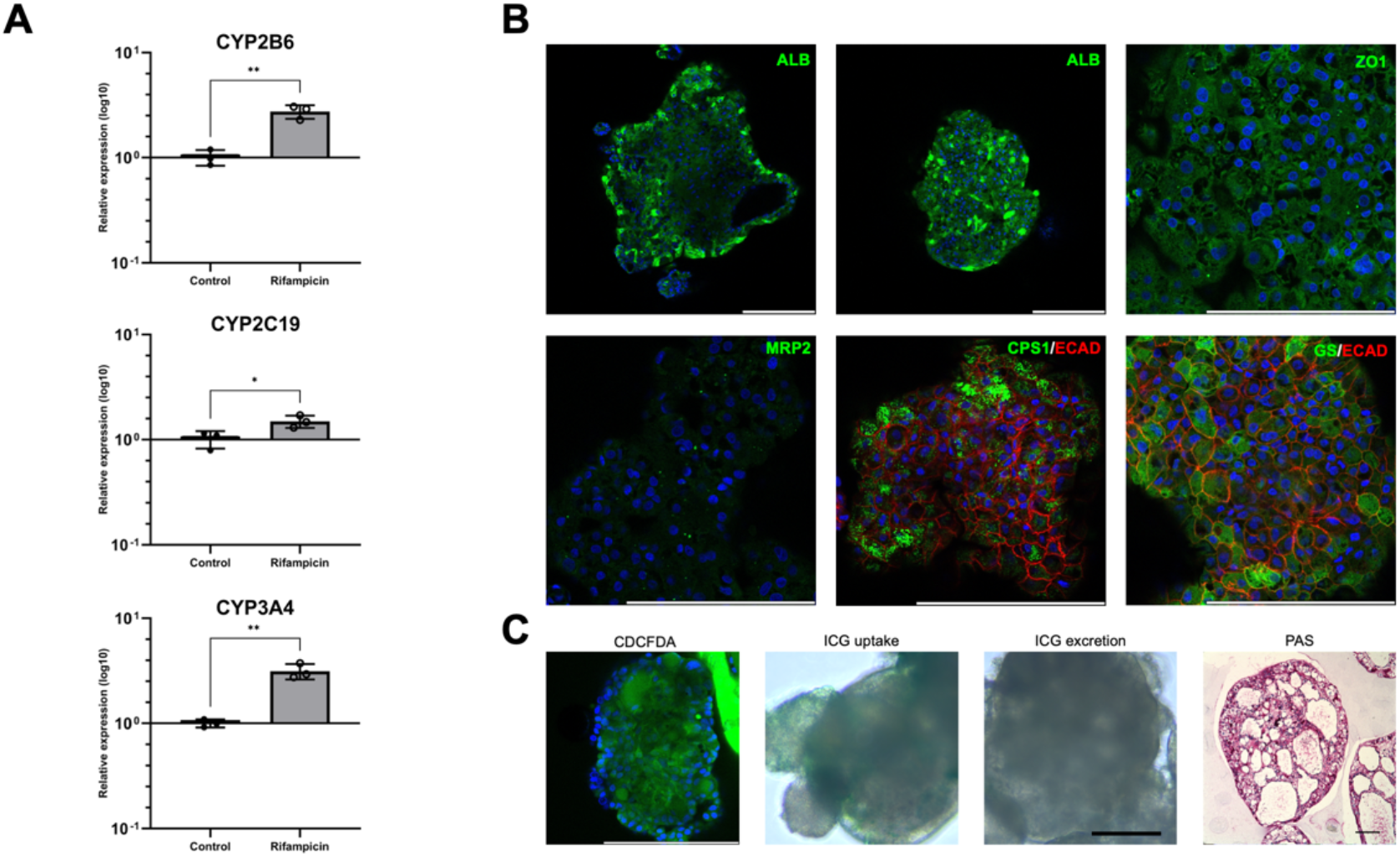
Liver functions of organoids. (A) RT-qPCR showing the extent of CYP2B6, 2C19 and 3A4 expressions in control and experimental organoids treated with 25 μM CYP inducer Rifampicin for 72 hours. (B) Immunofluorescent staining for ALB, AAT, ECAD, CPS1, GS, ZO1 and MRP2 expressions. (C) CDCFDA efflux assay showing polarization of hepatocytes, ICG uptake in 1 hour and excretion in 4 hours and PAS staining in paraffin section of organoids. Results are shown as mean ± SEM (n=3). *p <0.05; **p <0.01; ***p <0.001 using 2-tailed student’s *t* test. Scale bar = 200 μm.

### Liver organoids show organ specific structure and presence of liver specific cell types

Using immunofluorescent staining for CD31 and CD146, we detected a cluster of endothelial cells located around the center of the organoid 48 hours after cell seeding. On day 14, these endothelial cells have proliferated and formed an extensive vascular network throughout the organoids. CD31^+^ vasculatures possessed a hollow lumen, and some of the cells that lined these lumen also co-expressed CD34 (Fig. 5A). (24). We also observed that the CD44^+^ mesenchymal population was first evenly dispersed throughout the organoids when on day 2, but then formed into a thin layers lining the ECAD^+^ hepatocytes layer on day 14 (Fig. 5A). This spatial organization was comparable with the *in vivo* localization of HSCs, which normally reside in proximity to hepatocytes inside the perisinusoidal space of Disse (6). β-catenin was found localized on cell membranes of the outer epithelial cell layer (Fig. 5B), suggesting their involvement in maintenance of epithelial integrity by forming β-catenin/ECAD complexes (25). Laminin is a component of the sinusoidal basement membrane and was detected in the organoids as thin sheets lining both outer and inner surfaces of epithelial layer, which is consistent with their *in vivo* location in space of Disse (26). HNF4A is a hepatic marker and SOX9 is an early marker expressed in both hepatoblasts and cholangiocytes. Double immunofluorescent staining for HNF4A and SOX9 revealed the HNF4A^+^ hepatic population containing some of hepatoblasts co-expressed SOX9 (Fig. 5B). The cells that were SOX9^+^ HNF4A^-^ lining the inner luminal spheres, were identified as early biliary cells (27).

**Fig. 5.**
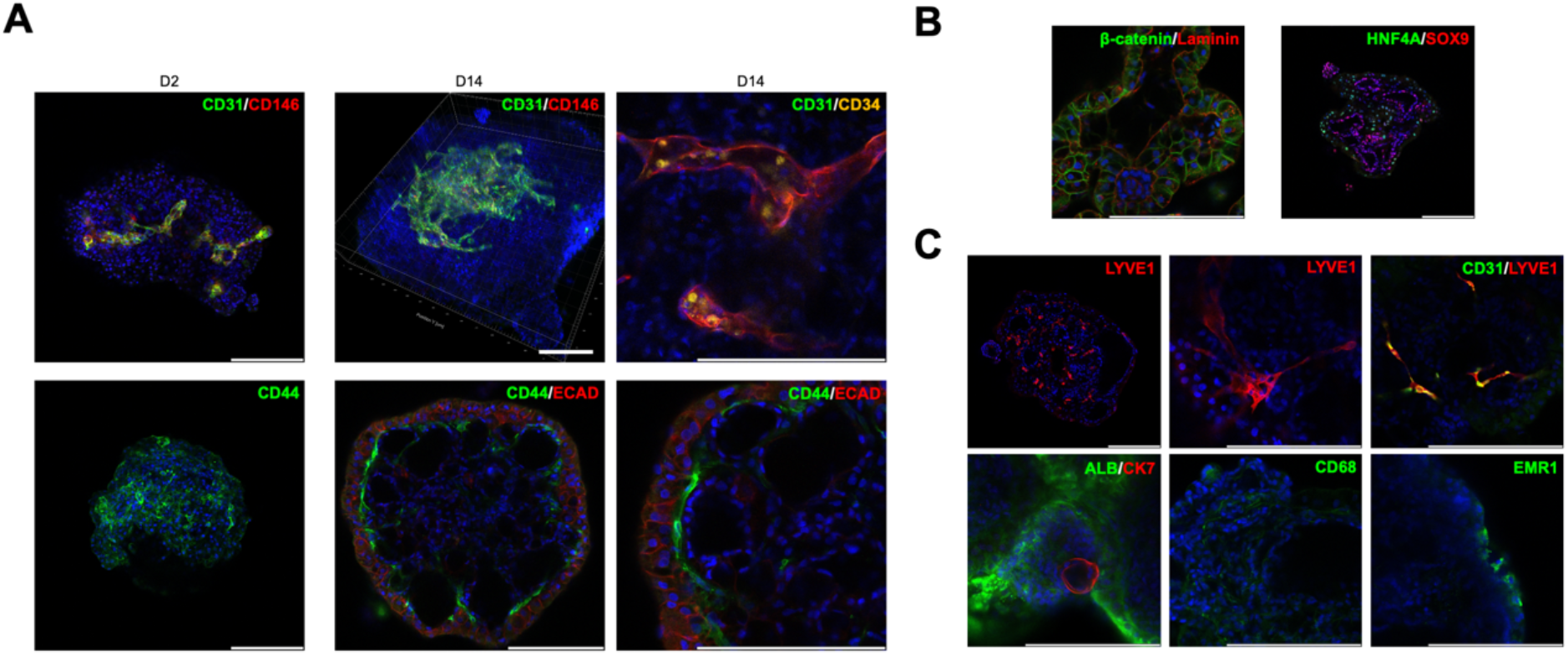
Spatial localization of PCs and NPCs inside liver organoids. (A) Immunofluorescent staining for CD31/CD146 in Day 2 and Day14 post-cultured organoids, CD31/CD34 in Day 14 organoids, CD44 in Day 2 and CD44/ECAD in Day 14 organoids. (B) Immunofluorescent staining for β-catenin (BCAT)/ Laminin and HNF4A/SOX9 in Day 14 organoids. (C) Immunofluorescent staining for LYVE1, LYVE1/CD31, ALB/CK7, CD68 and EMR1 in Day 14 organoids. Scale bar = 200 μm.

To investigate whether iPSC-derived ECs had the potential to differentiate into liver specific cell type in organoid culture, we conducted immunofluorescent staining with liver sinusoidal endothelial cells (LSECs) marker LYVE1 antibody (28). LYVE1^+^ cells were detected throughout liver organoids, with some forming tube-like structure and co-expressed CD31 (Fig. 5C). We also investigated the expression of mature cholangiocyte marker CK7 and macrophage marker CD68 and EMR1. ALB^-^ CK7^+^ circular luminal structure that resembled in vivo cholangiocytes (29), as well as CD68^+^ and EMR1^+^ macrophage populations were identified (Fig. 5C) (30). These finding revealed the developmental and differentiation potentials of our organoids to produce cells type normally found in liver NPC population.

### Liver fibrogenesis can be induced in liver organoids by thioacetamide (TAA)

We assessed the potential of organoids to become a liver fibrosis model by subjecting our liver organoids to 7-days of repeat-dose of TAA. The TAA treated organoids showed no obvious change in the size and morphology, but more dead cells present detaching from the organoids (Fig. 6A). ATP assay revealed a decrease in cell viability in organoids treated with 10 mM (68.3 ± 28.8% of control viability) and 25 mM TAA (30.3 ± 9.7% of control viability) (Fig. 6A). We confirmed that there was CYP2E1 activity in the liver organoids because bioactivation of TAA requires CYP2E1 which turns the TAA into hepatotoxic metabolites (31). 10 mM TAA did not cause reduction in liver function, but organoids treated with 25 mM TAA secreted significantly less albumin (33.3 ± 6.7% of control secretion) (Fig. 6A). 25 mM TAA induced significant up-regulation in fibrillar ECM including COL1A1 and COL3A1 (Fig. 6B). In addition, increase expression of ECM regulators MMP2 and TIMP1, HSC activation marker PDGFRB, fibrosis marker LOXL2 (32,33), pro-inflammatory cytokines IL6 and TNFA in our liver organoids, which allude to the early onset of inflammation and fibrosis (Fig. 6B). (34).

**Fig. 6.**
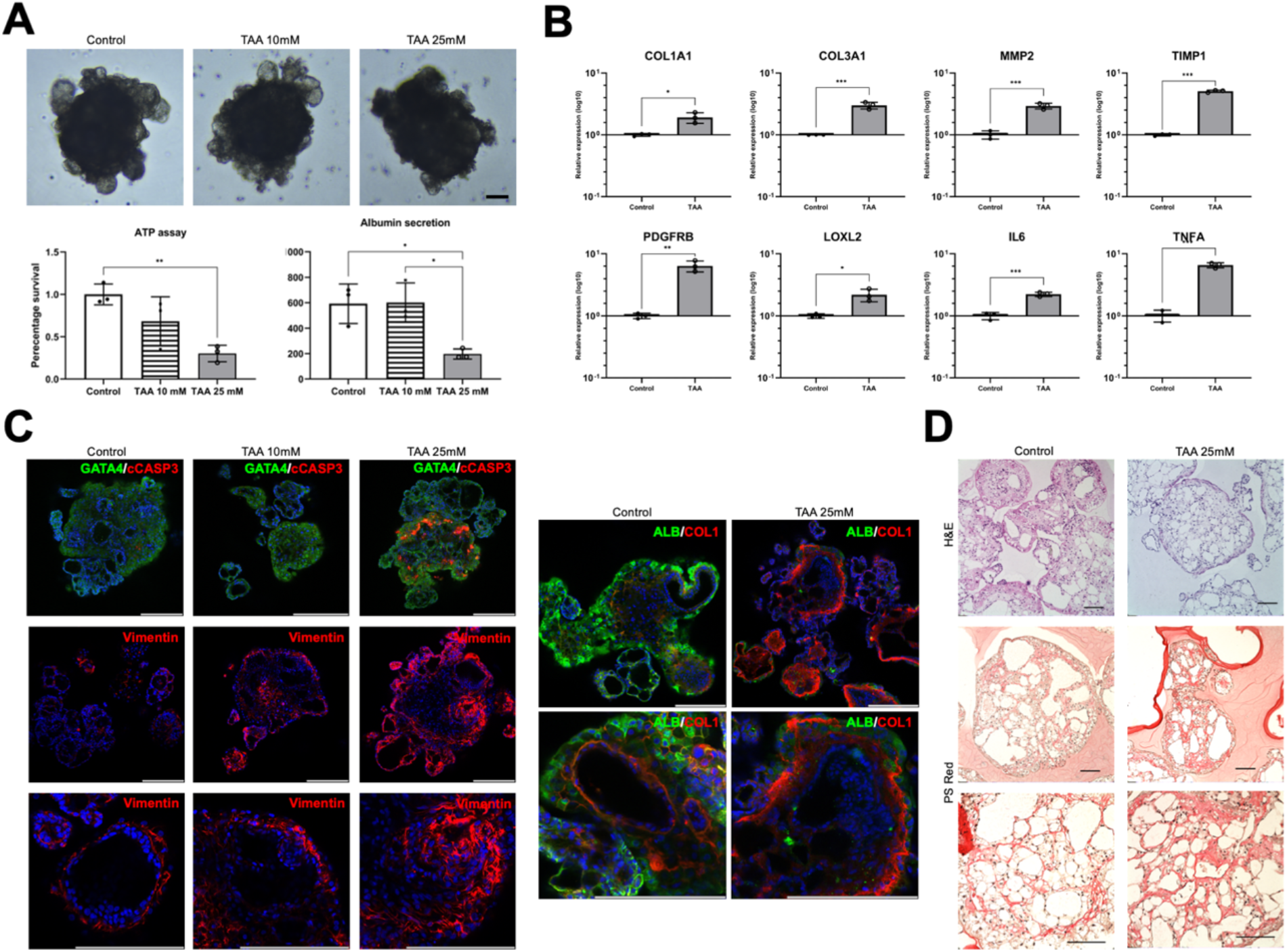
TAA induced liver fibrosis in liver organoids. (A) Morphology, cell viability and albumin secretion of control organoids and organoids treated with 10 or 25 mM TAA. (n=3) RT-qPCR showing COL1A1 and COL3A1, MMP2, TIMP1, PDGFRB, LOXL2, IL6 and TNFA expressions in different organoids. (C) Immunofluorescent staining of cleaved caspase 3 (cCASP3), vimentin and ALB/COL1 of different groups. (D) H&E and Picro Sirius Red (PSRed) stained paraffin sections of different groups. Results are shown as mean ± SEM (n=3). *p <0.05; **p <0.01; ***p <0.001 using 2-tailed student’s *t* test. Scale bar = 200 μm.

TAA-induced apoptosis and HSC activation was further confirmed with immune-fluorescent staining. At 25 mM TAA exposure, the liver organoids showed strong expression of apoptosis marker cleaved caspase 3 (cCASP3) (Fig. 6C). There was also increased vimentin staining in organoids treated with 10 and 25 mM TAA indicating an increase in HSC proliferation (35). In control organoids, the outer hepatocyte layer expressed ALB, while COL1 was detected as a thin layer surrounding biliary cells that lined the inner lumens, and on the inner cell surface of hepatocyte layer (Fig. 6C). This is consistent with *in vivo* liver histology as fibrillar collagens are normally found around the portal triads and in the perisinusoidal space (36). Organoids treated with 25 mM TAA lost most of their ability to express ALB and produced excess collagens in the perisinusoidal space reminiscent of liver fibrosis. The collagen deposits were also revealed by Picro Sirius Red (PS Red) staining (Fig. 6D).

### Free fatty acids (FFA) treatment induces steatosis and fibrosis in liver organoids

Liver organoids were treated with 200, 400 and 800 μM FFA for 7 days to induce steatosis. The organoids exhibited dose-dependent accumulation of neutral lipid droplets stained as revealed by Nile Red dye (Fig. 7A). Organoids treated with 800 μM FFA turned darker in appearance following lipid loading. FFA treatment did not cause significant cell death, but there was a significant decrease in albumin secretion at 200 and 800 μM (70.1 ± 7.7% and 49.9 ± 4.2% of control). Confocal image of the organoid surface displayed a ballooning of hepatocytes accompanied with a loss of ALB expression (Fig. 7A) (37). There was significant induction of COL1A1, COL3A1, TIMP1, PDGFRB, LOXL2 and TNFA gene expressions (Fig. 7B). This suggests that our organoids were displaying both inflammatory and fibrotic responses, which are more complex than simple steatosis in NAFLD. The results were further confirmed by immunofluorescent staining, which revealed an increased in vimentin expressing HSC population and increase in COL1 deposits (Fig. 7C). Double immunostaining for CD44 and COL3 revealed increased migration of mesenchymal cells and their secretion of COL3 that resembled the *in vivo* formation of fibrous septa, which was also displayed in PS Red stained paraffin sections (Fig. 7D).

**Fig. 7.**
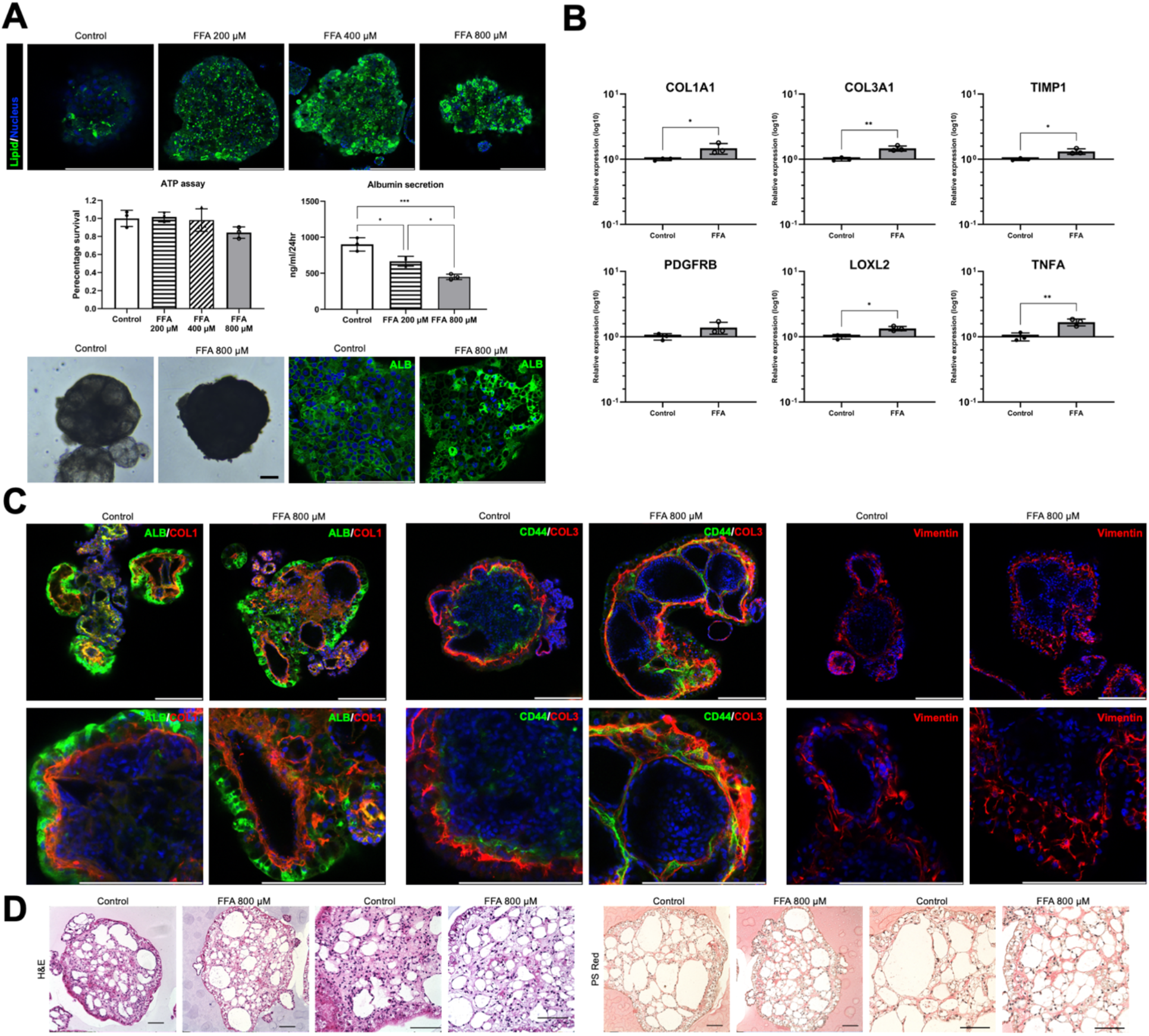
FFA induced liver steatosis and fibrosis in liver organoids. (A) Lipid droplets stained by Nile Red dye, cell morphology, cell viability, albumin secretion and ALB immunofluorescent staining of control organoids and organoids treated with different concentration of FFA. (n=3) (B) RT-qPCR showing COL1A1 and COL3A1, TIMP1, PDGFRB, LOXL2 and TNFA expression in different organoids. (C) Immunofluorescent staining for vimentin, ALB/COL1 and CD44/COL3 in different organoids. (D) H&E and Picro Sirius Red (PSRed) stained of paraffin sections of different organoids. Results are shown as mean ± SEM (n=3). *p <0.05; **p <0.01; ***p <0.001 using 2-tailed student’s *t* test. Scale bar = 200 μm.

## DISCUSSION

iPSC-derived liver organoid offers a promising alternative to 2D monolayer culture of hepatic cell lines and primary cells as an *in vitro* model, because of the unlimited availability of iPSCs and superior liver functions. However, current organoids lack of cellular complexity and great intra and inter batch variability have impeded their application in disease modeling and high throughput drug screening. In this study, we have described a protocol for generating liver organoids composed of HBs, MSCs, ECs, HSCs and entirely derived from a single hiPSC line. These different hepatic cell types were able to self-organize to create a highly vascularized liver organoid that displayed liver-specific functions and architecture post-14 days culture. Compared with organoids solely composed of iPSC-derived HB, organoids formed with iPSC-derived HBs and NPCs enhanced HBs maturation into hepatocytes. This was validated by significant increase in albumin secretion and expression of mature hepatocyte markers, CYP enzymes and liver function genes. These findings authenticate the supportive roles of NPCs in hepatic organoid differentiation through paracrine communication and direct cell interactions.

Our liver organoid model contains functional hepatocytes that were organized as a polarized epithelial layer around the periphery of the organoid. A layer of HSCs lined the inner surface of this hepatocyte layer, which histologically mimics the localization of HSCs in the space of Disse (6). Thin sheets of laminin and fibrillar collagen were found lining the hepatocytes which were comparable with the basement-like ECM found *in vivo*. (26) Our organoid were also highly vascularized, where the endothelial core formed during the early organization was found to have sprouted and extended into a network. The extensive vascularization explains why there was an absence of necrotic cores inside organoids despite of their large size, as vascularization facilitates the transport and diffusion of oxygen and nutrients into the centre of organoids. An abundance of LSECs were also found, which suggests the presence of a suitable micro-environment for endothelial progenitor cells or mesodermal cells to differentiate into liver specific ECs inside organoids. The complex cellular composition of our liver organoid is further confirmed by the presence of CK7^+^ luminal structures and EMR1^+^ macrophages.

Our liver organoids possess liver-mimetic functions and histotypic organization which make them promising for modeling compound-induced fibrosis and NAFLD, by recapitulating many of the complex events interactions between PCs and NPCs. Hepatotoxicity and declined hepatic functions caused by TAA reflected the CYP2E1 enzyme activities in hepatocytes (31), and hepatocytes had the ability to take up and store FFA as intracellular lipid droplets in a dose-dependent manner (38). FFA also hindered liver functions and caused hepatocyte ballooning after 7-day treatment, although without significant increase in cell death (5). The injured hepatocytes then activated the HSCs and macrophages, causing them to over-express genes associated with inflammatory cytokines, activation of HSCs and fibrosis, with increased HSC proliferation. Activated HSCs secreted excess fibrillar collagens that were observed to be distributed in a fibrous septa-like pattern.

Our ability to create liver organoids of consistent size, cellular composition and high albumin secretion allow individual organoid to be directly compared. This minimizes the intra-batch variations and number of organoids needed when applied in high-throughput screening for potential drug candidates. Further investigation will be needed to assess the efficiency of our organoids for screening potential candidates of anti-fibrotic and anti-steatotic drugs. These organoids also show potential in detecting drugs that cause liver injury, as well as in engineering of transplantable liver tissues.

## Abbreviations

EC: endothelial cell
FFA: free fatty acid
HB: hepatoblast
HLC: hepatocyte-like cell
HSC: hepatic stellate cell
hiPSC: human induced pluripotent stem cell
LSEC: liver sinusoidal endothelial cell
MSC: mesenchymal stem cell
NAFLD: non-alcoholic fatty liver disease
NASH: non-alcoholic steatohepatitis
NPC: non-parenchymal cell
PC: parenchymal cell
TAA: thioacetamide

## Author’s contribution

H.Y.T designed and performed the experiments, analyzed the data and wrote the manuscript.

P.H.Y.L and K.K.H.L consulted on study design. K.K.H.L supervised and reviewed the manuscript.

