## Supplementary Information for "Generation of liver organoids from human induced pluripotent stem cells as liver fibrosis and steatosis models"

### Supplementary Methods

#### Cell culture

Bone marrow-derived mesenchymal stem cells (BMMSCs) isolated from a young Caucasian male were cultured on gelatin coated dishes in MEM alpha supplemented with 10% FBS, 1% PS and 2 mM L-Glutamine, and passaged every 5 to 6 days at the ratio of 1:3.

Human umbilical vein endothelial cells (HUVECs) were cultured on gelatin coated dishes and passaged routinely at the ratio of 1:4 in M200 medium supplemented with 2% Large Vessel Endothelial Supplement (LVES) and 0.2% Gentamicin/Amphotericin).

Immortalized hepatic stellate cell (HSC) line LX-2 was maintained in DMEM-High Glucose, 10% FBS and 1% PS and passaged every week at the ratio of 1:10.

#### Differentiation of iPSCs

*Hepatocyte-like cells* iPSCs were differentiated into definite endoderm, hepatic progenitor and hepatocytes according Sullivan's group protocol with modifications (1). In brief, iPSCs were seeded at a density of 30,000 cells per cm$^2$ before culturing in endodermal differentiation medium (RPMI1640 medium, 2% B27 supplement without insulin, CHIR99021 4 μM) for 24 hours, then in fresh endodermal differentiation medium without CHIR99021 for 24 hours. Medium was replaced with hepatic specification medium (knockout-DMEM, 20% knockout serum replacement (KOSR), 2mM L-glutamine, 1% non-essential amino acids, 0.1mM β-mercaptoethanol and 1% DMSO) from day 2 to day 7 with medium change every 48 hours. On day 7, medium was replaced with hepatic maturation medium (L-15, 10% fetal bovine serum, 10% tryptose phosphate broth, 2mM L-glutamine, 1% Insulin-Transferrin-Selenium, 50μg/ml L-ascorbic acid, 10μM Hydrocortisone 21-hemisuccinate, supplemented with 100nM dexamethasone and 100nM Dihexa) and medium was changed every 48 hours until day 21.

*Mesenchymal stem cells* iPSCs were differentiated into MSCs according to Chen's group protocol with modifications (2). When iPSC culture reached confluence, medium was replaced with fresh MSC differentiation medium (DMEM:Ham's F-12 1:1 mixture, 20% KOSR, 1mM L-Glutamine, 1% NEAA, 10 μM SB431542) every day for 10 days. On day 10, cells were reseeded onto 0.1% gelatin coated dishes at the density of 4 x 10$^4$ cells per cm$^2$ in MSC culture medium (MEM alpha, 10% FBS, 1% P/S, 2mM L-Glutamine) and passaged every 4-5 days. Only passages 6 – 10 were used in downstream experiments.

*Endothelial cells* iPSCs were differentiated according to Patsch's group protocol with modification (3). iPSCs were at the density of 8,000 cells per cm$^2$ and cultured in mesoderm differentiation medium (DMEM/F12, 1% B-27 supplement (minus insulin), 2 mM LGlutamine, 8 μM CHIR99021, 25 ng/ml human recombinant BMP4 (Peprotech)) when cells reached 50% confluence with medium change every 48 hours for 4 days. Medium was then replaced with EGM-2 (Lonza) without FBS, supplemented with 100 ng/ml human recombinant VEGF165 (Peprotech) and 2 μM forskolin (Meryer), with medium change every day for 3 days before FACS for CD31$^+$ cells using BD FACSAria II cell sorter. CD31$^+$ cells cultured in Endothelial Cell Medium (Sciencell)and passaged every 5-6 days. Only passages 2 – 4 were used in downstream experiments.

*Hepatic stellate cells* iPSCs were differentiated into hepatic stellate cells according to Coll's group protocol (4). In brief, iPSCs were first seeded at the density of 9 x $10^4$ cells per $cm^2$. When cells reached 50% confluency, medium was replaced with HSC differentiation medium (57% DMEM-Low Glucose, 40% MCDB-201 Water, 0.25x linoleic acid-bovine serum albumin, 0.25x insulin-transferrin-selenium, 1% penicillin streptomycin, 100 μM Lascorbic acid, 2.5 μM dexamethasone, 50 μM 2-mercaptoethanol) for 12 days with medium change every 48 hours. Different growth factors and chemicals were supplemented into the differentiation medium at different time: 4 μM CHIR99021 and 20 ng/ml rh BMP4 on day 1 to 6, 20 ng/ml rh FGF1 and 20 ng/ml rh FGF3 on day 5 to 8, 5 μM retinol and 100 μM palmitic acid on day 6 to 12. On day 12, cells were reseeded and cultured in DMEM GlutaMax supplemented with 10% FBS, 1% PS, 5 μM retinol and 100 μM palmitic acid up to 4 passages.

**RNA extraction and real-time quantitative polymerase chain reaction (RT-qPCR)**

Total RNA was extracted using RNAiso Plus reagent (TaKaRa) with or with a Direct-zol RNA microprep kit (Zymo Research) according to the manufacturer's instructions. RNA quantified using Nanodrop 2000 Spectrophotometer (ThermoFisher Scientific). Reverse transcription was performed with 2μg RNA from each sample using PrimeScript RT Master Mix (TAKARA). RT-qPCR was performed using SYBR Premix Ex Taq (Tli RNaseH Plus) (TAKARA) on QuantStudio 7 Flex Real Time PCR System (Applied Biosystems).

**Liver function test**

*Urea colourimetric assay* Culture media was collected stored at -80°C until use. Colourimetric assay was performed using Urea Nitrogen (BUN) Test (Stanbio) according to manufacturer's protocol with 450 μl reaction volume for each sample.

*Period Acid and Schiff (PAS) Staining* Cells fixed with 4% PFA for 10 mins, or rehydrated paraffin sections of organoids were incubated in 1% periodic acid for 15 mins, followed by 15 mins incubation in Schiff's solution.

*Oil Red O Staining* Oil Red O stock solution was prepared by dissolving 0.5g Oil Red O in 100ml isopropanol, then was filtered using 0.45 μm syringe filter. Prior to use, Oil Red O stock solution was added to dd $H_2O$ at 3:2 ratio, mixed well and incubated for 10 mins before being filtered using 0.45 μm filter again. Cells were fixed in 4% PFA for 30 mins, incubated in 60% isopropanol for 5 mins followed by 15 mins incubation in Oil Red O solution.

*CDCFDA Staining* Organoids were incubated in 5 μM 5-(and-6)-Carboxy-2',7'-Dichlorofluorescein Diacetate (CDCFDA) in PBS for 20 mins in dark, then washed twice in ice-cold PBS and observed using IX83 inverted fluorescent microscope (Olympus).

*Nile Red Staining* Organoids were fixed in 4% PFA for 1 hour then incubated in 1mg/ml Nile Red solution for 2 hours before observation under confocal microscopy. Nuclei were counterstained with 5μg/ml DAPI.

**iPSC-MSC multilineage differentiation**

*Osteogenic and Adipogenic differentiation* Confluent iPSC-MSCs were cultured in osteogenic differentiation medium (alpha MEM, 10% FBS, 1% P/S, 2 mM GlutaMax, 10 mM β-glycerophosphate, 100 μg/ml L-Ascorbic acid, 100 nM dexamethasone) or adipogenic differentiation medium (alpha MEM, 10% FBS, 1% P/S, 2 mM GlutaMax, 10 μg/ml insulin, 500 μM IBMX, 200 μM indomethacin, 1 μM dexamethasone) with medium change thrice a week for 28 days. Differentiated osteoblasts were stained with Alizatin Red S dye for calcium deposits and BCIP/NBT solution for alkaline phosphatase activity. Differentiated adipocytes were stained for oil droplets with Oil Red O.

*Chondrogenic differentiation* iPSC-MSCs were trypsinized and 2.5 x 105 cells were pelletd in a 15-ml. Chondrogenic differentiation medium (alpha MEM, 1% P/S, 2 mM GlutaMax, 1 mM sodium pyruvate, 40 μg/ml L-Proline, 1% ITS, 100 μg/ml L-Ascorbic acid, 100 nM dexamethasone, 10 ng/ml TGFB3) was added on top without disturbing the cell pellet. Pellets were cultured in tube for 28 days with medium change thrice a week. Tube cap was loosely screwed for gas exchange. Pellets were harvasted, embedded in paraffin blocks and sectioned for Alcian Blue staining.

**Tube formation assay**

48- well plates were first coated with Matrigel (50 μl per cm2) for 1 hour in 37°C, before plating 2.5 x $10^4$ iPSC-ECs per $cm^2$ in EGM-2. After 16 hours in 37°C, 5% CO2 incubater, the formed tubes were incubated in 2 μM Calcein AM in PBS for 20 mins in dark, washed with PBS thrice and observed under fluorescent microscopy.

**HSC activation by TGFB1**

iPSC-HSCs were starved for 24 hours in serum free medium, then treated with 5 or 10 ng/ml TGFβ1 for 72 hours with medium change every day.

**Differential trypsinization of iPSC-HBs**

On Day 7 of differentiation, iPSC-HBs were washed with PBS twice and treated for 5 mins in 1mM EDTA for 5 min, then 5 mins in TryPLE Express solution at 37°C. Undifferentiated cells detached form culture dish and were gently removed. Remaining attached cells were incubated in fresh TryPLE Express for 5 mins, neutralized 10% FBS and collected for organoid formation.

**Free fatty acid-BSA conjugation**

1M oleic acid or 500 mM palmitic acid dissolved in 100% ethanol was diluted in 37.5mM NaOH and heated at 70°C for 30 mins to generate 20 mM sodium oleate or sodium palmitate. Sodium oleate or sodium palmitate was further diluted to 5mM in 5% fatty acid-free BSA (Solarbio) dissolved in William's E medium and heated in 37°C oven for 1 hour with shaking. Result free fatty acid-BSA conjugate (FFA-BSA) had free fatty acid to BSA ratio of 6:1. BSA control was prepared by heating 5% BSA with NaOH solution without FFA.

**Supplementary Tables**

**Table S1. List of antibodies used in this study.**

| Antigen | Dilution | Company |
|---|---|---|
| AAT | 1:100 | Santa Cruz |
| ALB | 1:100 | Santa Cruz |
| ARG1 | 1:100 | Santa Cruz |
| cCASP3 | 1:200 | Cell signaling |
| CD31 | 1:200 | BD Pharminogen |
| CD34 | 1:200 | Abcam |
| CD44 | 1:100 | BD Pharminogen |
| CD68 | 1:100 | Santa Cruz |
| CK18 | 1:100 | Santa Cruz |
| CK19 | 1:100 | Santa Cruz |
| CK7 | 1:200 | ABclonal |
| COL1 | 1:200 | Abcam |
| COL3 | 1:200 | Abcam |
| CPS1 | 1:100 | Santa Cruz |
| CYP3A4 | 1:100 | Santa Cruz |
| ECAD | 1:200 | Cell signalling |
| EMR1 | 1:100 | Santa Cruz |
| EPCAM | 1:100 | Santa Cruz |
| FOXA2 | 1:100 | Santa Cruz |
| GS | 1:100 | Santa Cruz |
| HNF4A | 1:100 | Santa Cruz |
| Ki67 | 1:200 | Cell signalling |
| Laminin | 1:200 | ABclonal |
| LYVE1 | 1:100 | R&D |
| MRP2 | 1:100 | Santa Cruz |
| PDGFRB | 1:100 | BD Pharminogen |
| SOX9 | 1:100 | Santa Cruz |
| TBX3 | 1:100 | Santa Cruz |
| TNFA | 1:200 | ABclonal |
| Vimentin | 1:200 | ABclonal |
| ZO1 | 1:100 | Santa Cruz |

| | | |
|---|---|---|
| β-catenin | 1:200 | ThermoFisher |

**Table S2. List of primers used in this study.**

| Names of genes | Primer sequences (5' to 3') | |
|---|---|---|
| GAPDH | Forward | TTGAGGTCAATGAAGGGGTC |
| | Reverse | GAAGGTGAAGGTCGGAGTCA |
| FOXA2 | Forward | TACGTGTTCATGCCGTTCAT |
| | Reverse | CGACTGGAGCAGCTACTATGC |
| GATA4 | Forward | CGACTTCTCAGAAGGCAGAGAGTG |
| | Reverse | CTTCATGTAGAGGCCGCAGGCATT |
| SOX17 | Forward | CGCACGGAATTTGAACAGTA |
| | Reverse | GGATCAGGGACCTGTCACAC |
| AFP | Forward | GCAGCCAAAGTGAAGAGG |
| | Reverse | TGTTGCTGCCTTTGTTTG |
| HNF4A | Forward | CCAAGTACATCCCAGCTTTC |
| | Reverse | TTGGCATCTGGGTCAAAG |
| EPCAM | Forward | ATAACCTGCTCTGAGCGAGTG |
| | Reverse | TGAAGTGCAGTCCGCAAACT |
| CK19 | Forward | ACCAAGTTTGAGACGGAACAG |
| | Reverse | CCCTCAGCGTACTGATTTCCT |
| SOX9 | Forward | AGCGAACGCACATCAAGAC |
| | Reverse | GCTGTAGTGTGGGAGGTTGAA |
| TBX3 | Forward | CCCGGTTCCACATTGTAAGAG |
| | Reverse | GTATGCAGTCACAGCGATGAAT |
| ALB | Forward | GGCACAATGAAGTGGGTAAC |
| | Reverse | AGGCAATCAACACCAAGG |
| AAT | Forward | ACTGTCAACTTCGGGGACAC |
| | Reverse | CATGCCTAAACGCTTCATCA |
| TAT | Forward | ACTGTGTTTGGAAACCTGCC |
| | Reverse | GCAGCCACTTGTCAGAATGA |
| CYP3A4 | Forward | AAGTCGCCTCGAAGATACACA |
| | Reverse | AAGGAGAGAACACTGCTCGTG |
| MRP2 | Forward | TCTCTCGATACTCTGTGGCAC |

| | | |
|---|---|---|
| | Reverse | CTGGAATCCGTAGGAGATGAAGA |
| HNF1B | Forward | CCCAGCAAATCTTGTACCAGGC |
| | Reverse | ACCTCAGTGACCAAGTTGGAGC |
| HNF6 | Forward | GAGGATGTGGAAGTGGCTGCAG |
| | Reverse | CTGTGAAGACCAACCTGGGCTT |
| OCT4 | Forward | CCTGAAGCAGAAGAGGATCACC |
| | Reverse | AAAGCGGCAGATGGTCGTTTGG |
| SOX2 | Forward | GGAAAACCAAGACGCTCATGAAGAAGG |
| | Reverse | GTTCATGTAGGTCTGCGAGCTGGTCAT |
| NANOG | Forward | CCTCCATGGATCTGCTTATTCAGGACA |
| | Reverse | CCTTCTGCGTCACACCATTGCTATTCT |
| CMYC | Forward | GCTGCTTAGACGCTGGATTT |
| | Reverse | CACCGAGTCGTAGTCGAGGT |
| ZEB1 | Forward | AGGCAGATGAAGCAGGATGT |
| | Reverse | CTCTTCAGGTGCCTCAGGAA |
| ZEB2 | Forward | CAAGCACCACCTTATCGAGC |
| | Reverse | TGTGATTCATGTGCTGCGAG |
| NCAD | Forward | AGGGATCAAAGCCTGGAACA |
| | Reverse | TTGGAGCCTGAGACACGATT |
| SNAI1 | Forward | TTACCTTCCAGCAGCCCTAC |
| | Reverse | GACAGAGTCCCAGATGAGCA |
| SNAI2 | Forward | CCCATGCCATTGAAGCTGAA |
| | Reverse | TTTCTAGACTGGGCATCGCA |
| CD29 | Forward | CAAAGGAACAGCAGAGAAGC |
| | Reverse | ATTGAGTAAGACAGGTCCATAAGG |
| CD44 | Forward | CAACAACACAAATGGCTGGT |
| | Reverse | CTGAGGTGTCTGTCTCTTTCATCT |
| CD73 | Forward | CAGTACCAGGGCACTATCTGG |
| | Reverse | AGTGGCCCCTTTGCTTTAAT |
| CD105 | Forward | CCACTAGCCAGGTCTCGAAG |
| | Reverse | GATGCAGGAAGACACTGCTG |
| CD34 | Forward | TCTGATCTCCATGGCTTCCT |
| | Reverse | ACTGAGGCAACAGCTCAACC |
| CD31 | Forward | ATTGCAGTGGTTATCATCGGAGTG |
| | Reverse | CTCGTTGTTGGAGTTCAGAAGTGG |
| VECAD | Forward | TCGTCATGGACCGAGGTT |

| | | |
|---|---|---|
| | Reverse | TCTACAATCCCTTGCAGTGTGA |
| FLT1 | Forward | CCTGCAAGATTCAGGCACCTATG |
| | Reverse | GTTTCGCAGGAGGTATGGTGCT |
| FLT4 | Forward | CTCTGCCTGGGACTCCTG |
| | Reverse | GGTGTCGATGACGTGTGACT |
| NKX2.5 | Forward | ATCCTAAACCTGGAGCAGCA |
| | Reverse | AGATCTTGACCTGCGTGGAC |
| vWF | Forward | CCTTGAATCCCAGTGACCCTGA |
| | Reverse | GGTTCCGAGATGTCCTCCACAT |
| eNOS | Forward | GCGGCTGCATGACATTGAG |
| | Reverse | GTCGCGGTAGAGATGGTCAAG |
| EphrinB2 | Forward | CTCCTCAACTGTGCCAAACCA |
| | Reverse | GGTTATCCAGGCCCTCCAAA |
| CoupTF2 | Forward | TGGTTCCAAACCAGTTTATTCTGTG |
| | Reverse | AAGTGCGTTTCCATCATCTTTGAG |
| NOTCH1 | Forward | CAGGCAATCCAGGACTATG |
| | Reverse | CAGGCGTGTTGTTCTCACAG |
| LYVE1 | Forward | ACTTCCATCTGGACCA |
| | Reverse | CCTTTTTGCTCACAAG |
| STAB2 | Forward | CCTGTGAAACCTGTGCTGAC |
| | Reverse | CCATCGCCATCTAGTCCACTG |
| ICAM1 | Forward | AGCGGCTGACGTGTGCAGTAAT |
| | Reverse | TCTGAGACCTCTGGCTTCGTCA |
| CD32B | Forward | AGTGCCCAGCATGGGCAGC |
| | Reverse | TGTCTCTTTCTGATGGCAAT |
| CD45 | Forward | CTTCAGTGGTCCCATTGTGGTG |
| | Reverse | CCACTTTGTTCTCGGCTTCCAG |
| FLK1 | Forward | CCTGTATGGAGGAGGAGGAA |
| | Reverse | CGGCTCTTTCGCTTACTGTT |
| DESMIN | Forward | AGGAACAGCAGGTCCAGGTA |
| | Reverse | AGAGCATCAATCTCGCAGGT |
| GFAP | Forward | GGATGGAGAGGTCATTAAGGA |
| | Reverse | GGTGAGTTTCTTGTTAGTTGGA |
| PDGFRB | Forward | CCCTTATCATCCTCATCATGC |
| | Reverse | CCTTCCATCGGATCTCGTAA |
| NCAM | Forward | AGGAGACAGAAACGAAGCCA |

| | | |
|---|---|---|
| | Reverse | GGTGTTGGAAATGCTCTGGT |
| HGF | Forward | CGCTGGGAGTACTGTGCAAT |
| | Reverse | CCCTGTAGCCTTCTCCTTGA |
| LRAT | Forward | TACTGCAGATATGGCACCCC |
| | Reverse | CCAAGACTGCTGAAGCAAGA |
| RELN | Forward | GTCTACCTTCCACTCTCCACCA |
| | Reverse | GTCCAGCATCACAAATCCCTCG |
| PCDH7 | Forward | GAGGAGTCAGAAACACCAAGCAG |
| | Reverse | TCAGGGCTACATCTGGAAGAGG |
| PDGFRA | Forward | AACCCTGCTGATGAAAGCAC |
| | Reverse | TCCTTTCTAGCATGGGGACA |
| ALCAM | Forward | CTTCTGCCTCTTGATCTCCG |
| | Reverse | AGGTACGTCAAGTCGGCAAG |
| CYGB | Forward | TCTATGCCAACTGCGAGGAC |
| | Reverse | TCCTCCATGTGCTTGAACTG |
| PPARG | Forward | GAGGAGAGTTACTTGGTCGT |
| | Reverse | CAACAGACAAATCACCATTCG |
| LOXL2 | Forward | GGAGAGGACATACAATACCAAAGTG |
| | Reverse | CCATGGAGAATGGCCAGTAG |
| aSMA | Forward | CCAGAGCCATTGTCACACAC |
| | Reverse | CAGCCAAGCACTGTCAGG |
| COL1 | Forward | GTGCGATGACGTGATCTGTGA |
| | Reverse | CGGTGGTTTCTTGGTCGGT |
| MMP1 | Forward | TTGTGGCCAGAAAACAGAAA |
| | Reverse | TTCGGGGAGAAGTGATGTTC |
| MMP3 | Forward | CAATTTCATGAGCAGCAACG |
| | Reverse | AGGGATTAATGGAGATGCCC |
| TIMP1 | Forward | CTGTTGTTGCTGTGGCTGAT |
| | Reverse | ACTTGGCCCTGATGACGAG |
| IL1B | Forward | ATGATGGCTTATTACAGTGGCAA |
| | Reverse | GTCGGAGATTCGTAGCTGGA |
| IL8 | Forward | ACTGAGAGTGATTGAGAGTGGAC |
| | Reverse | AACCCTCTGCACCCAGTTTTC |
| CEBPB | Forward | GCCCTCGCAGGTCAAGAGCA |
| | Reverse | TTGAACAAGTTCCGCAGGGTG |
| CK7 | Forward | TGTGGATGCTGCCTACATGAGC |

| | | |
|---|---|---|
| | Reverse | AGCACCACAGATGTGTCGGAGA |
| CD68 | Forward | CGAGCATCATTCTTTCACCAGCT |
| | Reverse | ATGAGAGGCAGCAAGATGGACC |
| CYP2B6 | Forward | GGCACACAGGCAAGTTTACA |
| | Reverse | CCAGCAAAGAAGAGCGAGAG |
| CYP2C9 | Forward | GGACAGAGACGACAAGCACA |
| | Reverse | TGGTGGGGAGAAGGTCAAT |
| CYP2C19 | Forward | ACTTGGAGCTGGGACAGAGA |
| | Reverse | CATCTGTGTAGGGCATGTGG |
| CYP7A1 | Forward | CACTTTGTCCACCTTTGATG |
| | Reverse | GCTGCTTTCATTGCTTCTG |
| G6PC | Forward | CTACAGCAACACTTCCGTGC |
| | Reverse | GTATACACCTGCTGTGCCCAT |
| COL3 | Forward | AGGACTGACCAAGATGGGAA |
| | Reverse | AGGGGAGCTGGCTACTTCTC |
| MMP2 | Forward | GGAAAGCCAGGATCCATTTT |
| | Reverse | ATGCCGCCTTTAACTGGAG |
| MMP9 | Forward | GCCACTACTGTGCCTTTGAGTC |
| | Reverse | CCCTCAGAGAATCGCCAGTACT |
| MMP13 | Forward | CCTTGATGCCATTACCAGTCTCC |
| | Reverse | AAACAGCTCCGCATCAACCTGC |
| TIMP2 | Forward | ACCCTCTGTGACTTCATCGTGC |
| | Reverse | GGAGATGTAGCACGGGATCATG |
| TIMP3 | Forward | TACCGAGGCTTCACCAAGATGC |
| | Reverse | CATCTTGCCATCATAGACGCGAC |
| IL10 | Forward | TCTCCGAGATGCCTTCAGCAGA |
| | Reverse | TCAGACAAGGCTTGGCAACCCA |
| IL12B | Forward | GACATTCTGCGTTCAGGTCCAG |
| | Reverse | CATTTTTGCGGCAGATGACCGTG |
| IL13 | Forward | ACGGTCATTGCTCTCACTTGCC |
| | Reverse | CTGTCAGGTTGATGCTCCATACC |
| IL6 | Forward | CCTGAACCTTCCAAAGATGGC |
| | Reverse | TTCACCAGGCAAGTCTCCTCA |
| SLCO1B1 | Forward | GCAATAGCAATGATTGGTCC |
| | Reverse | CCAACCCATCGAGAATCAG |
| TGFB1 | Forward | TACCTGAACCCGTGTTGCTCTC |

| | | |
|---|---|---|
| | Reverse | GTTGCTGAGGTATCGCCAGGAA |
| TNFA | Forward | CTCTTCTGCCTGCTGCACTTTG |
| | Reverse | ATGGGCTACAGGCTTGTCACTC |
| CCL2 | Forward | AGAATCACCAGCAGCAAGTGTCC |
| | Reverse | TCCTGAACCCACTTCTGCTTGG |
| CCL5 | Forward | CCTGCTGCTTTGCCTACATTGC |
| | Reverse | ACACACTTGGCGGTTCTTTCGG |

### Supplementary Figures

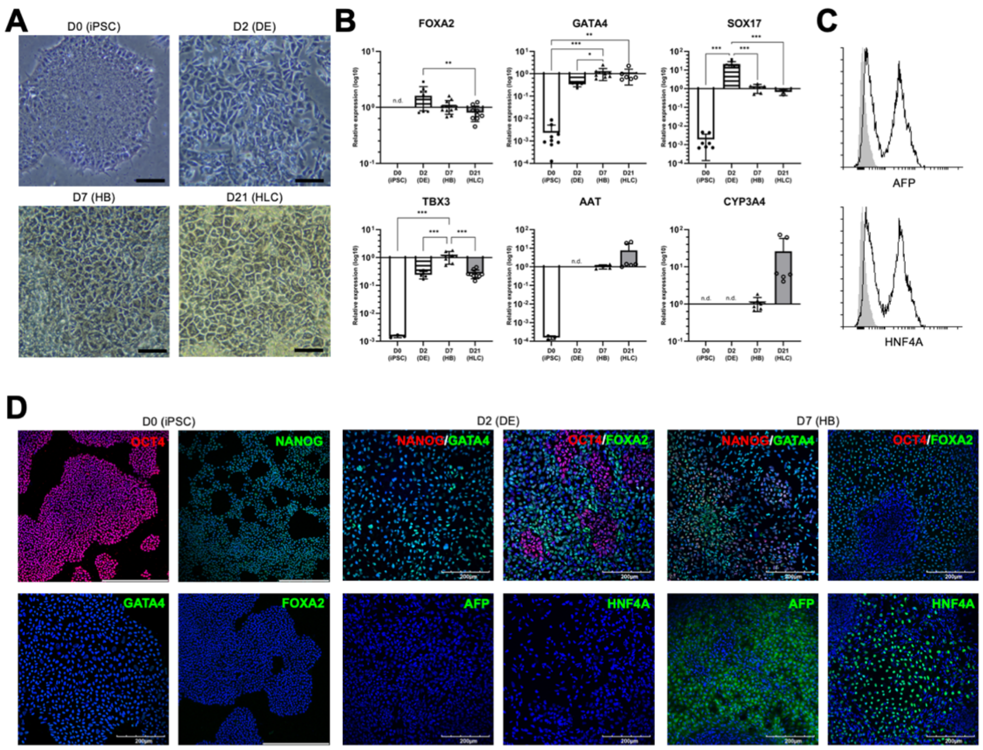


**Fig. S1. Hepatic differentiation of iPSCs.** (A) Morphology of iPSCs, Day (D)2 definite endoderm (DE), D7 hepatoblast (HB) and D21 hepatocyte-like cell (HLC). Scale bar = 50 µm. (B) Gene expression of definitive endoderm markers (FOXA2, SOX17, GATA4) and hepatocyte markers (TBX3, AAT, CYP3A4) in D2 DE, D7 HB and D21 HLC assessed by RT-qPCR using GAPDH for internal control. (C) Representative flow cytometry analysis of hepatoblast markers AFP and HNF4A in D7 HBs. IgG control displayed in grey. (D) Representative immunofluorescent staining images of markers for pluripotency markers (OCT4, NANOG) definitive endoderm (FOXA2, GATA 4) and hepatic marker (AFP, HNF4A) in iPSCs, D2 and D7 cells. Scale bar = 200 µm. Results are shown as mean ± SEM (n≥3). *p <0.05; **p <0.01; ***p <0.001 using 2-tailed student's *t* test.

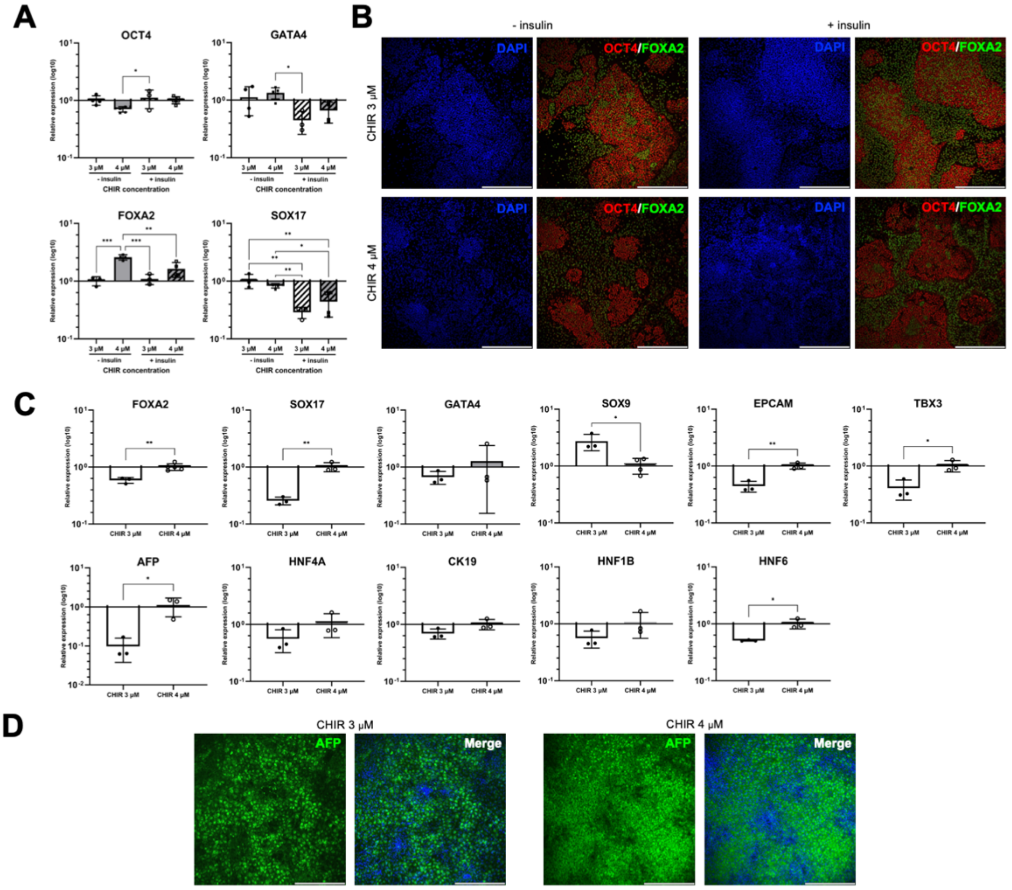


**Fig. S2. Optimization of endoderm and hepatoblast differentiation.** (A) Summary of different differentiation conditions. (B) Gene expression of pluripotency markers (OCT4, NANOG), definitive endoderm markers (FOXA2, SOX17, GATA4) in D2 DEs (CHIR 3 µM without insulin expression = 1) (C) Immunofluorescent staining images of FOXA2/OCT and NANOG/GATA4 in DEs. (D) Expression of endoderm and hepatoblast marker of D7 HBs treated with different CHIR concentrations at endoderm differentiation step. (CHIR 4 µM expression = 1) (E) Immunofluorescent staining images of AFP and HNF4A in HBs. (F) Flow cytometric analysis of endoderm and hepatoblast markers. (n>=3) Results are shown as mean ± SEM (n≥3). *p <0.05; **p <0.01; ***p <0.001 using 2-tailed student's *t* test. Scale bar = 200 µm.

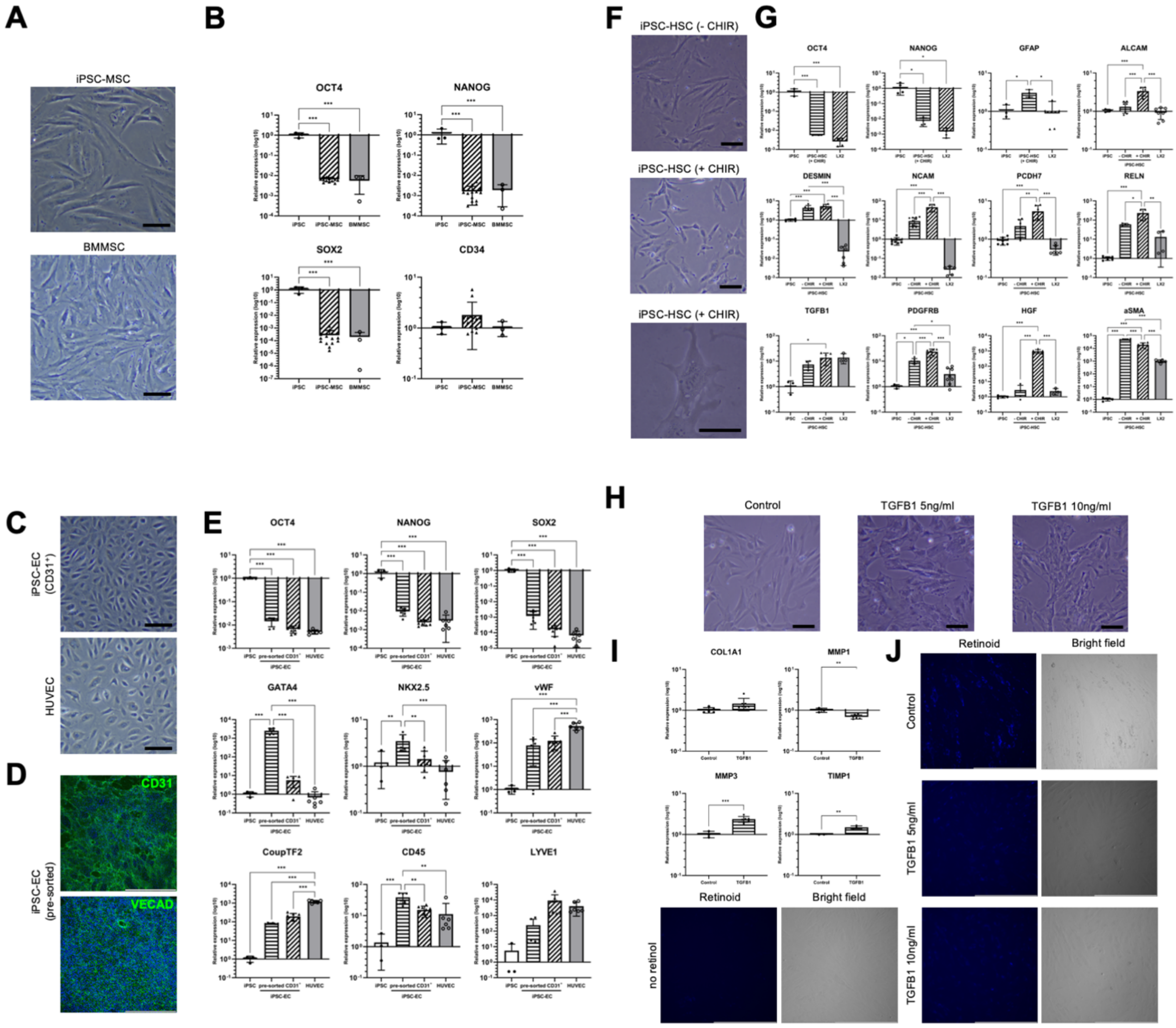


**Fig. S3. NPC differentiation of iPSCs.** (A) Morphology of iPSC-derived MSCs. They displayed elongated fibroblast-like morphology similar to bone marrow-derived MSCs (BMMSCs). Scale bar = 50 µm. (B) Expression of pluripotency markers (OCT4, NANOG, SOX2) and haematopoeitic marker (CD34) in iPSC-MSCs. (C) Morphology of iPSC-derived ECs. iPSC-ECs showed typical cobblestone endothelial morphology. Scale bar = 50 µm. (D) Immunofluorescent staining of EC markers CD31 and VECAD in pre-sorted iPSC-ECs. Scale bar = 200 µm. (E) Expression of pluripotency markers (OCT4, NANOG, SOX2), mesoderm markers (NKX2.5, GATA4) and EC markers (vWF, CoupTF2, CD45, LYVE1) by RT-qPCR. (F) Morphology of iPSC-derived HSCs differentiated with or without CHIR99021 for first 4 days. iPSC-HSCs differentiated with CHIR99021 exhibited a more star-like and less fibroblast-like morphology compared to those without using CHIR99021. Scale bar = 50 µm. (G) Expression of pluripotency markers (OCT4, NANOG) and HSC marker GFAP in iPSC-HSCs (with CHIR), and expression of ALCAM, DESMIN, NCAM, PCDH7, RELN, TGFB1) iPSC-HSCs with or without CHIR by RT-qPCR. (H) Morphology of TGFB1 treated iPSC-HSCs. Scale bar = 200 µm. (I) Expression of activation markers (COL1, MMP1, MMP3, TIMP1) in iPSC-HSCs. (J) Auto-fluorescent retinoid droplets in TGFB1 treated iPSC-HSCs. iPSC-HSCs showed a dose-dependent loss of intracellular retinoid droplets when treated with TGFB1. Results are shown as mean ± SEM (n≥3). *p <0.05; **p <0.01; ***p <0.001 using 2-tailed student's *t* test.

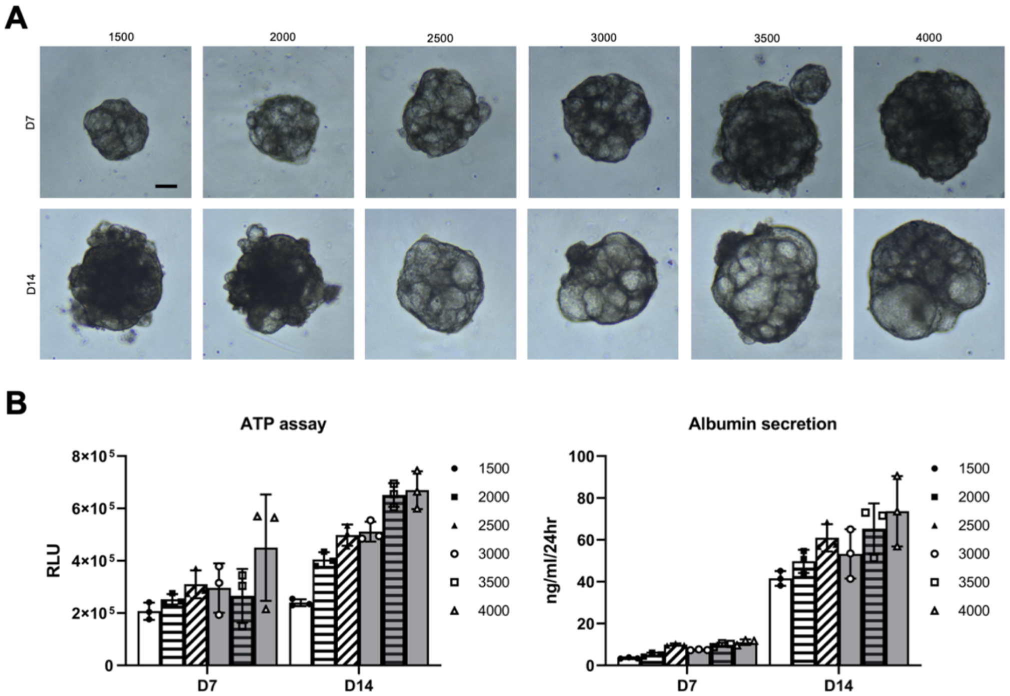


**Fig. S4. Effect of initial cell seeding number for iPSC-HB only organoid formation.** (A) Morphology of organoids formed with only iPSC-derived HBs at different seeding number on day 7 and 14. Scale bar = 200 μm. (B) Cell viability measured by ATP bioluminescence assay and albumin secretion measured by ELISA of organoids on day 7 and 14 at different seeding number. Results are shown as mean ± SEM (n=3).

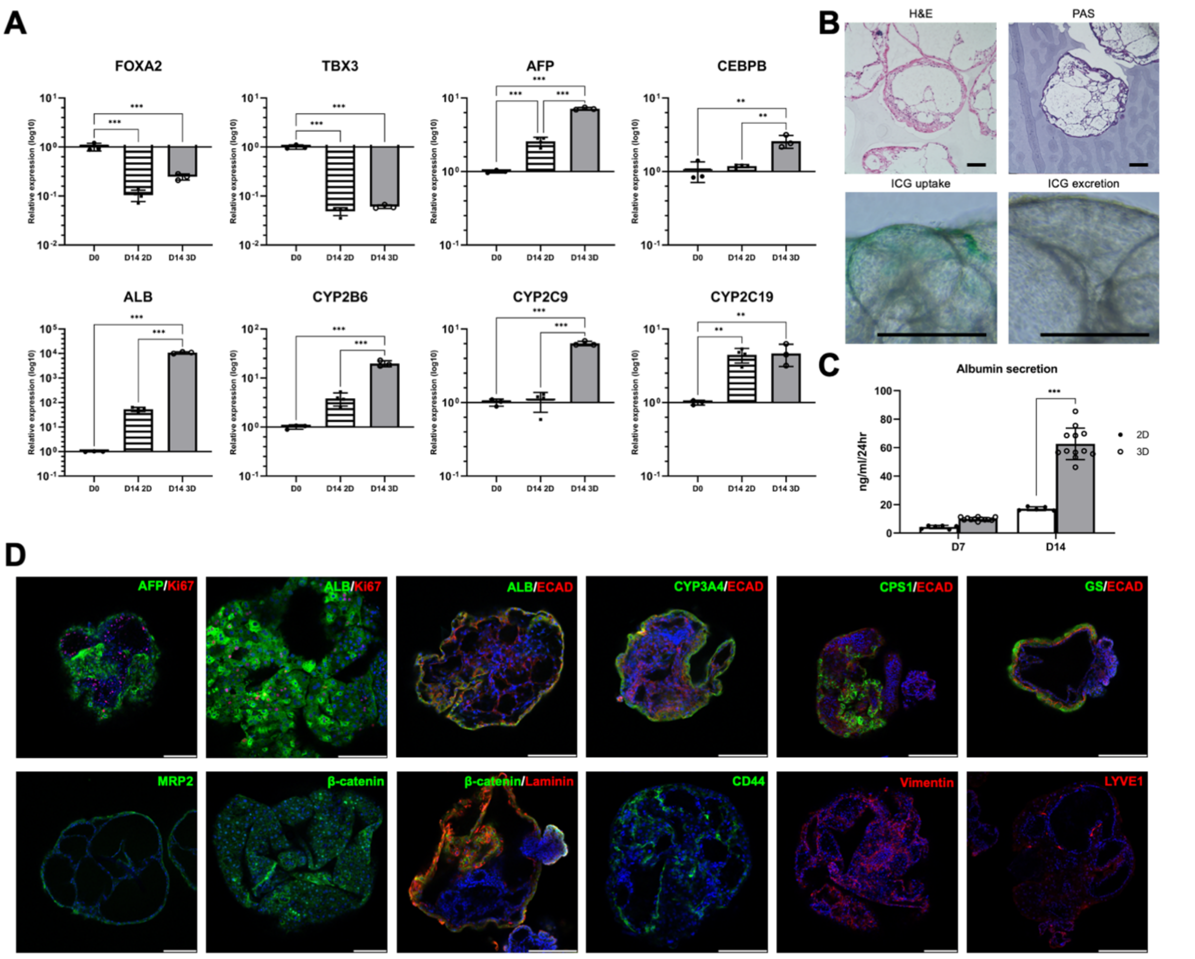


**Fig. S5. Characterization of iPSC-HB only organoid.** (A) Gene expression of endoderm marker (FOXA2), hepatoblast marker (TBX3, AFP), hepatocyte marker (CEBPB, ALB) and CYP enzymes (CYP2B6, 2C9, 2C19) in D0 freshly trypsinized iPSC-derived HBs, D14 2D culture and organoid culture of iPSC-HBs. D0 HB expression = 1. (B) H&E and PAS staining of organoid paraffin sections, indocyanine green uptake and excretion of organoids. (C) Albumin secretion of 2D culture and organoid culture of iPSC-HBs on day 7 and day 14. (D) Immunofluorescent staining of hepatocyte markers (ALB, ARG1, CYP3A4), epithelial marker (ECAD), hepatoblast markers (AFP, EPCAM, CK19), proliferation marker (Ki67), laminin and β-catenin. Results are shown as mean ± SEM (n≥3). *p <0.05; **p <0.01; ***p <0.001 using 2-tailed student's *t* test. Scale bar: 200 μm.

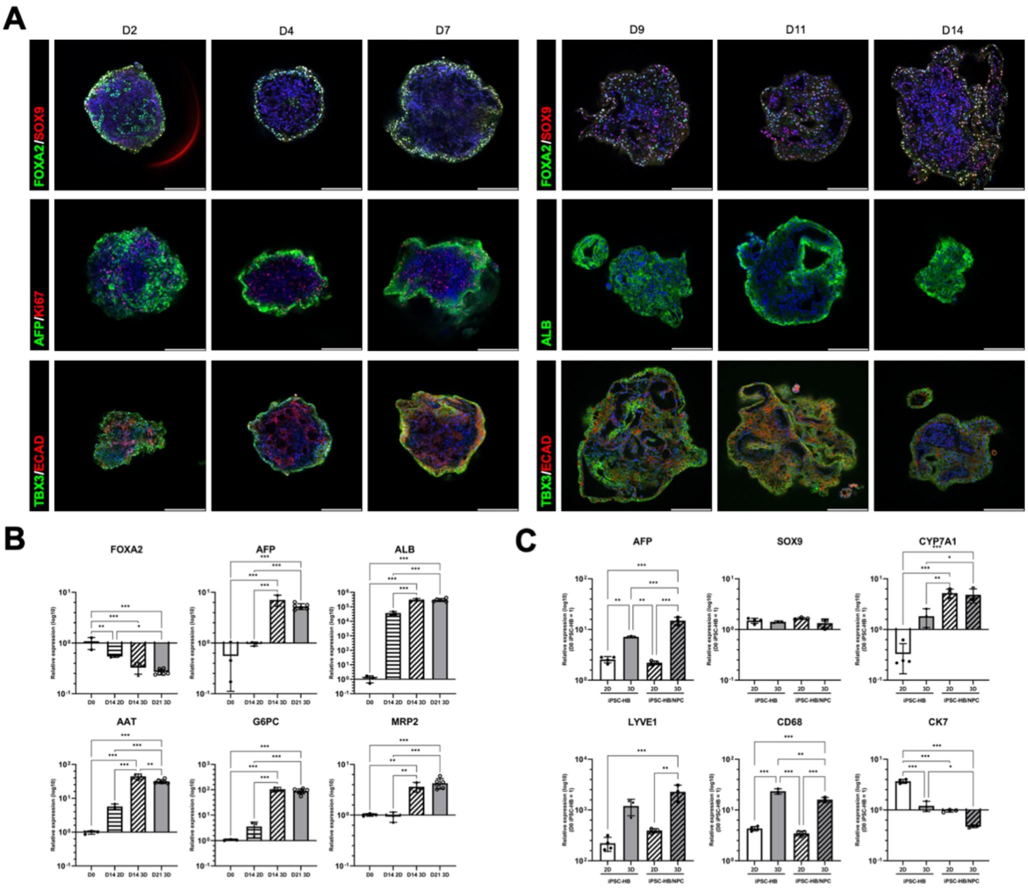


**Fig. S6. Characterization of iPSC-HB/NPC organoid.** (A) Immunofluorescent staining of FOXA2/SOX9, AFP/Ki67 and TBX3/ECAD in iPSC-HB/NPC organoids. Scale bar: 200 µm. (B) Gene expression of endoderm marker (FOXA2) and hepatocyte markers (AFP, ALB, AAT, G6PC, MRP2) in D0 freshly trypsinized iPSC-HB/NPC, D14 iPSC-HB/NPC 2D culture, D14 iPSC-HB/NPC organoids and D21 iPSC-HB/NPC organoids by RT-qPCR. Results are shown as mean ± SEM (n≥3). *p <0.05; **p <0.01; ***p <0.001 using 2-tailed student's *t* test. (C) Gene expression of hepatoblast marker (AFP, SOX9), hepatocyte markers (CYP7A1), LSEC marker (LYVE1), macrophage marker (CD68) and cholangiocyte marker (CK7) in D14 2D and 3D cultures of iPSC-derived HBs, D14 2D and 3D co-cultures of iPSC-derived HB and NPC by RT-qPCR as fold change to D0 trypsinized iPSC-HBs. Results are shown as mean ± SEM (n≥3). *p <0.05; **p <0.01; ***p <0.001 using 2-tailed student's *t* test.
